## Supplementary Figures for "Daily electric field treatment improves functional outcomes after thoracic contusion spinal cord injury in rats"

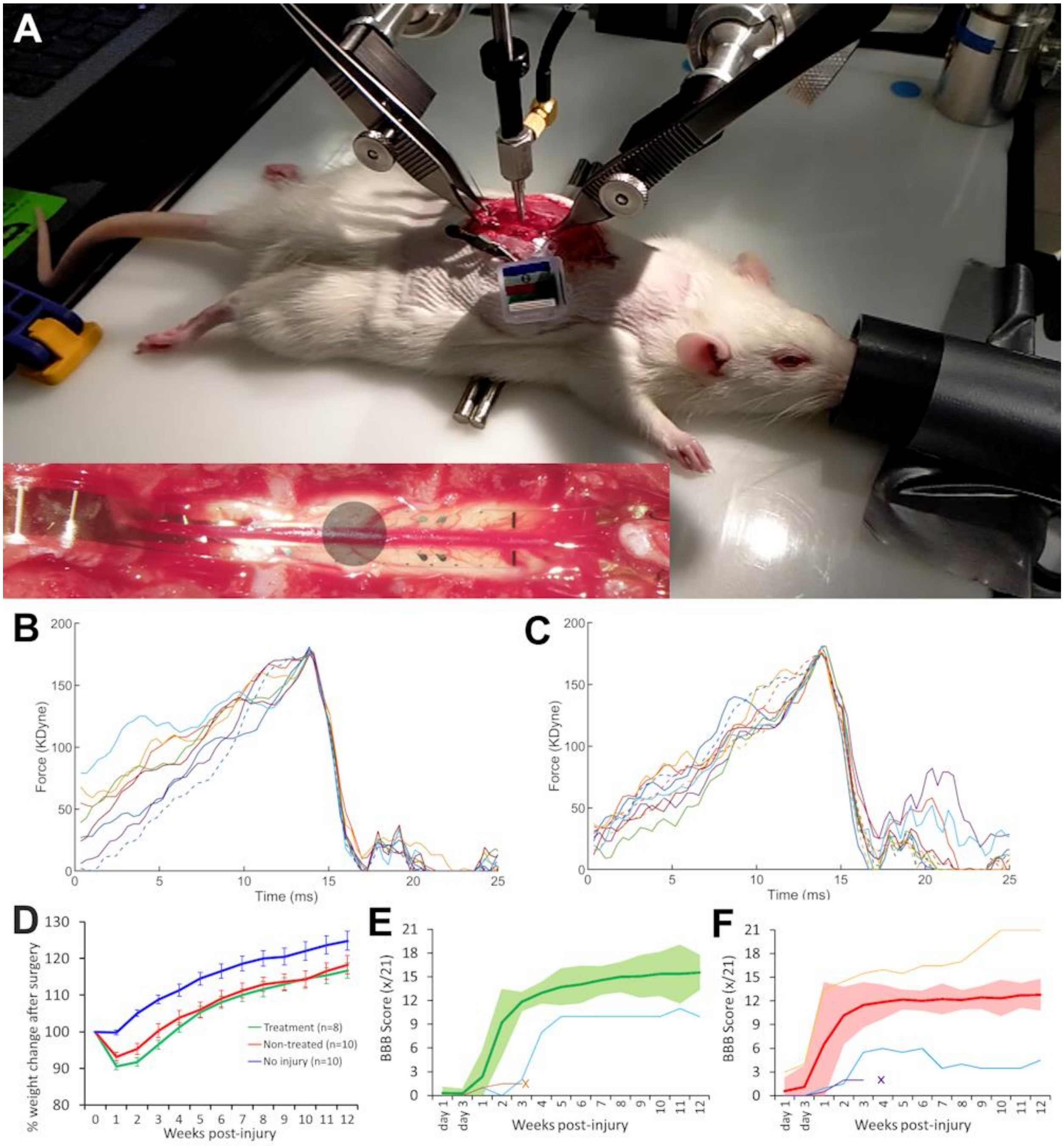


**Supplementary Figure 1:** Groups of treated and non-treated rats received a 175 kilodyne impact injury to spinal segment L1/L2 using an Infinite Horizons Impactor. (**A**) After subdural insertion of the implant, rats were placed on the impactor stage and held in position via hemostats clamped to the T9 and T13 spinal processes adjacent to the laminectomy. A 2.5 stainless steel impactor tip was used to deliver a 175 kilodyne impact to the boundary between spinal segments L1/L2 (directly below the T11 spinal process), which was centred between two sets of stimulation electrodes (shown in inset). Force vs time profiles of the impactions are shown for the (**B**) treated group and (**C**) non-treated group. (**D**) In both the treated and non-treated groups rat's weight dropped immediately after surgery and then steadily increased. (**E**) Two rats in the treated group were excluded as their injury progression was two standard deviations outside the group mean. (**F**) Three rats in the non-treated group were excluded as their injury progression fell two standard deviations outside the group mean. Two implants from excluded rats were explanted intact at weeks 3 and 4 post-surgery and electrochemically tested (time of explantation indicated by crosses).

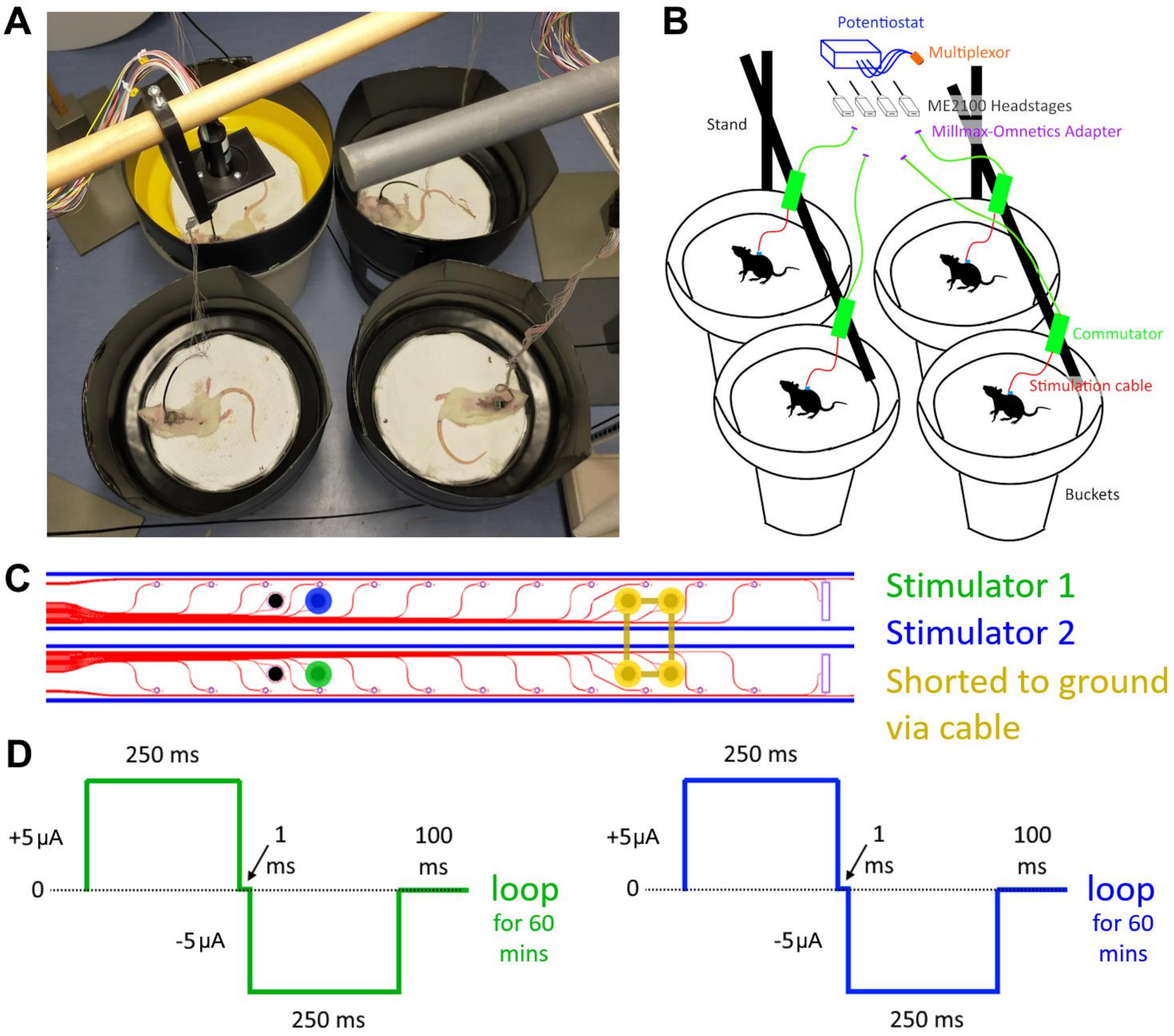


**Supplementary Figure 2:** Rats in the treated group received a daily one-hour EF treatment. (**A**) Up to four animals at a time were placed in comfortable towel-lined buckets and gently restrained while their implanted backpacks were connected to an overhanging stimulation cable. (**B**) The cables were connected to commutators, and each plugged into HS32 head stages and a ME2100 Multi Chanel Systems electrophysiology system for delivering stimulation. To test the impedance of electrodes, the cables exiting the commutators were instead plugged into a homemade multiplexor adapter and potentiostat. (**C**) Current-controlled ES was delivered via two stimulators in each headstage directed at stimulation electrodes on each arm of the implant (blue and green circles). The stimulation cables shorted the four electrodes on the caudal end to the ME2100 ground channel (yellow circles). (**D**) Each stimulator (green and blue) delivered a 1-hour electric field treatment consisting of looped ±5µA 250ms biphasic pulses with 100ms recharge periods across the injury site.

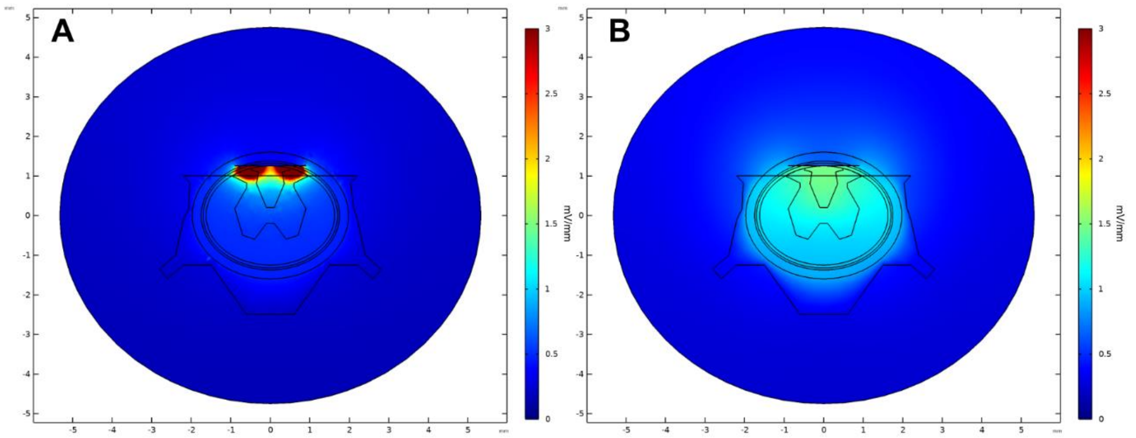


**Supplementary Figure 3:** Cross-sectional (coronal) perspective of FEM-based simulation of the medium sized rat's spinal cord showing longitudinal electric field strength at (**A**) the level of the electrodes, and (**B**) equidistant between the electrodes at the center of the estimated spinal lesion**.** The electric field is more uniform and extends to the ventral outline of the cord at the midpoint between the electrodes, compared to its distribution directly at the electrode positions.

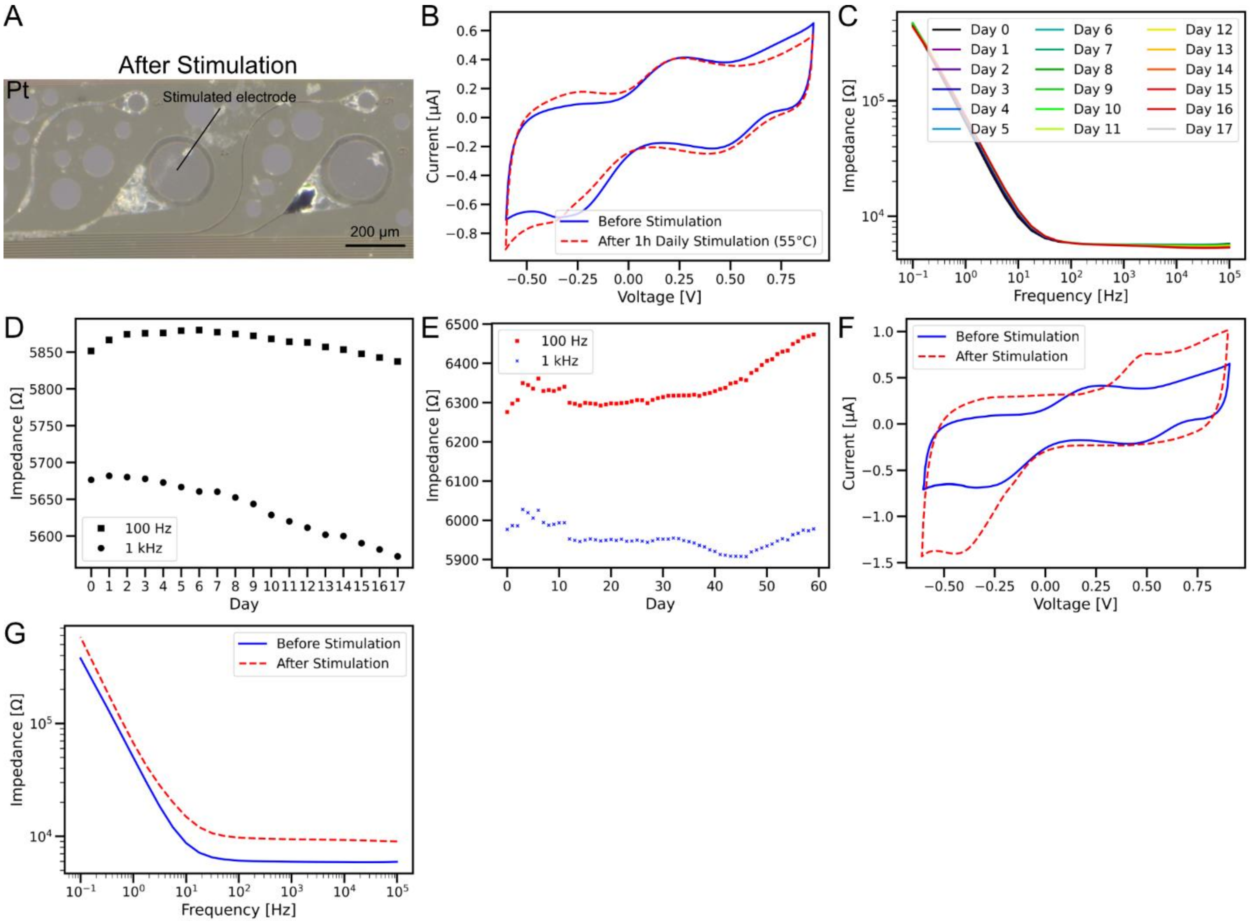


**Supplementary Figure 4:** The stability of Pt and SIROF stimulation electrodes was tested in 1x phosphate buffered saline using the same implant design but with fenestrations in the polyimide. (**A**) To test a Pt electrode, the SIROF coating was omitted. After 90 hours of continuous stimulation (5 µA, 250 ms, 2 Hz), the stimulated Pt electrode delaminated and dissolved, affecting neighboring recording and stimulation electrodes, which also delaminated and dissolved. We believe that over the course of the experiment first the stimulation electrodes dissolved. Then the Pt connection line was in contact with PBS and dissolved as well, causing a cross connection to neighboring channels. The *in vivo* treatment was simulated for 18 days at 55°C (equivalent to 62 days at 37°C according to the "10-degree rule") and for 60 days at 37°C. The same implant body was used for both temperatures, but different stimulation electrodes were tested. (**B, C, D**) After 18 days at 55°C with daily 1-hour stimulation, the SIROF electrode remained stable. (**E**) Similarly, after 60 days at 37°C with daily 1-hour stimulation, which is equivalent to the *in vivo* stimulation duration, the SIROF electrode remained stable. (**F, G**) Although the four shorted SIROF counter electrodes used at both temperatures showed changes in their CV and EIS, they remained functional.

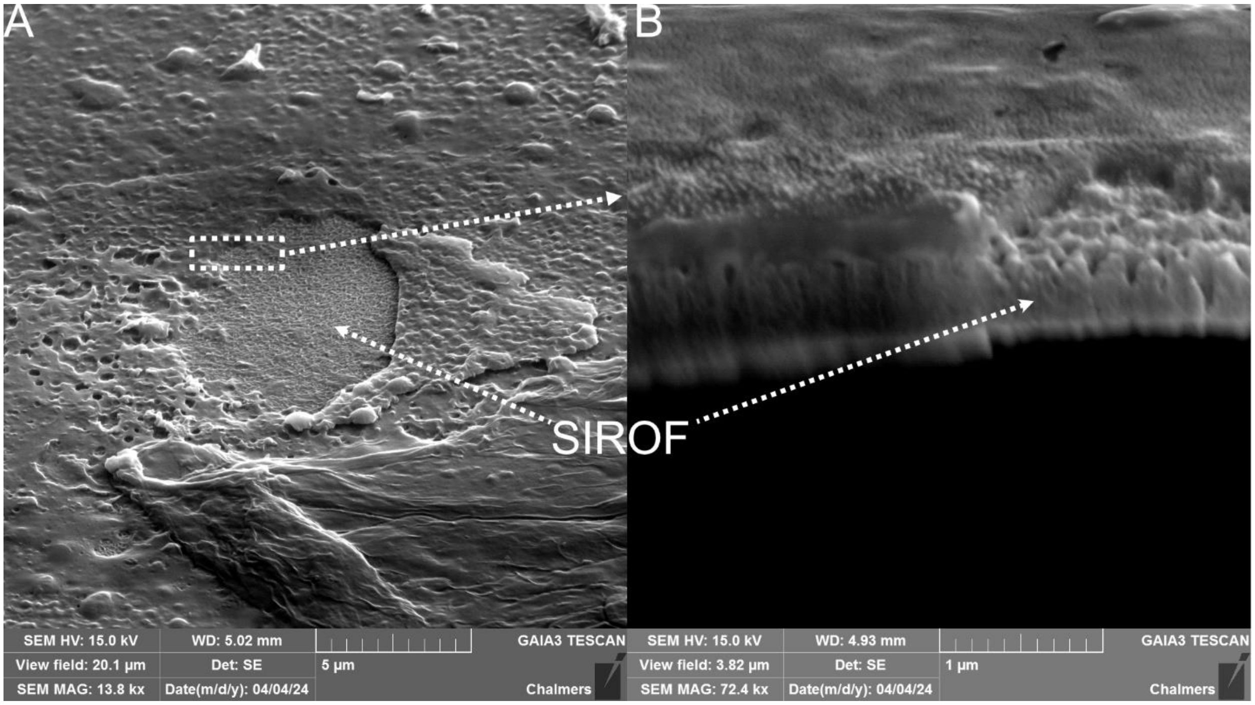


**Supplementary Figure 5:** (**A**) High resolution images of a CE SIROF electrode shorted during the *in vitro* stimulation test. (**B**) FIB-SEM of the SIROF shows a deposited layer on top of the SIROF.

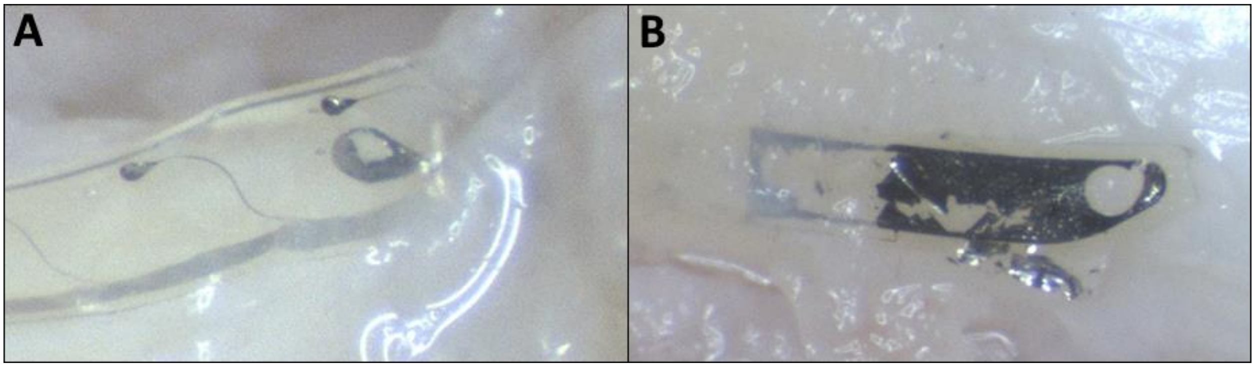


**Supplementary Figure 6:** During explantation of the device at 12 weeks, we observed that the larger electrode surfaces including the (**A**) stimulation electrodes and (**B**) large ground electrodes (used for recordings, not part of this study), delaminated from the polyimide and/or stuck to the surface of the spinal cord.

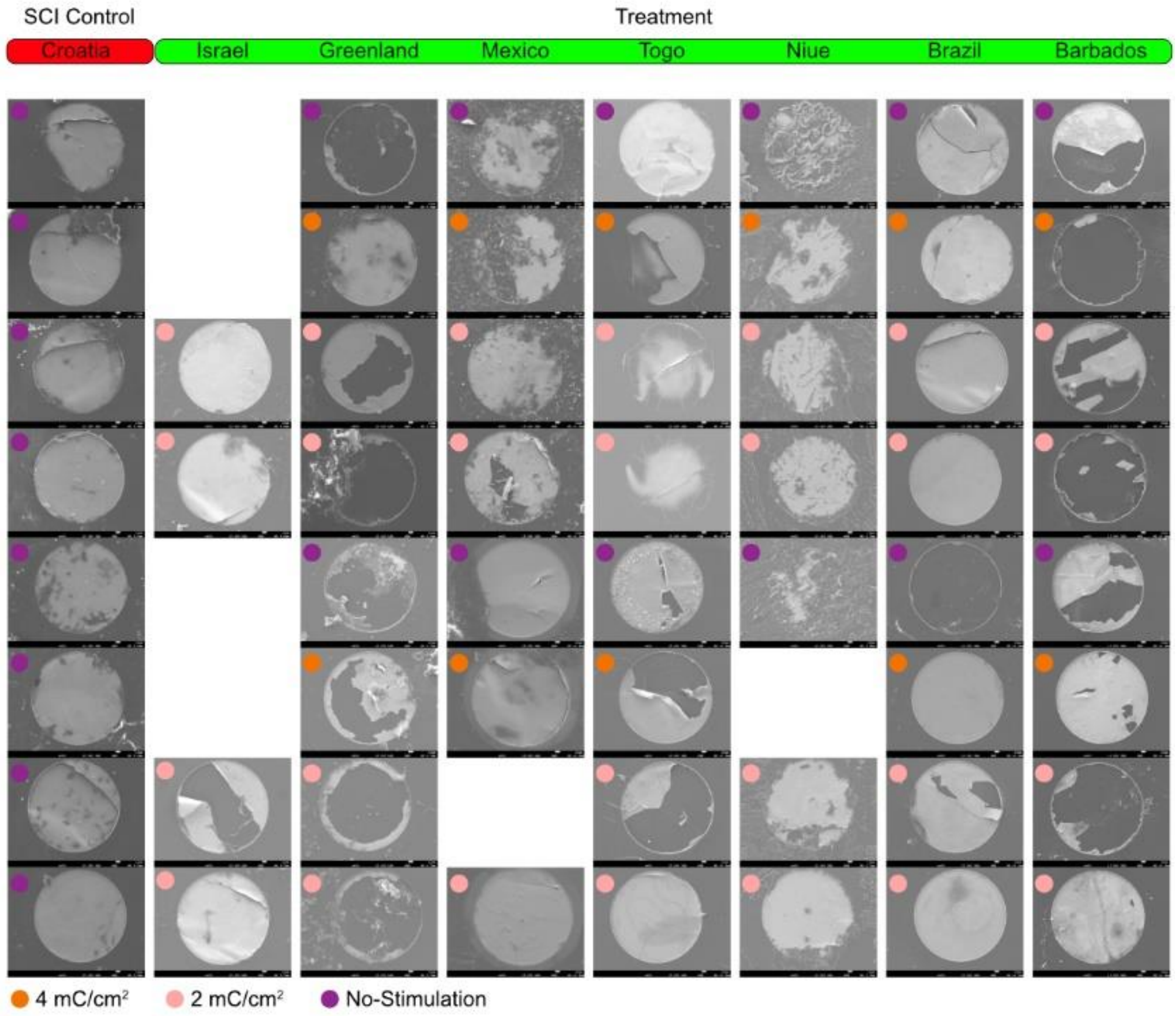

**Supplementary Figure 7**. SEM images of explanted devices.

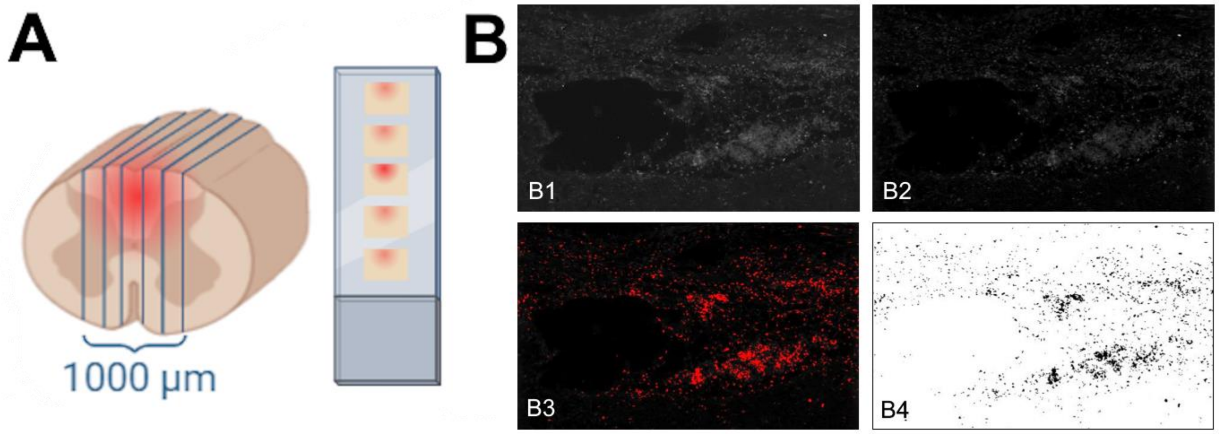


**C**

| Antibody | Host | Target | Isotype | Supplier | Cat # | Dilution |
|---|---|---|---|---|---|---|
| **Anti-GFAP** | Mouse | Astrocytes (indicates lesion area) | IgG1 | Sigma | G3893 | 1:500 |
| **Anti-GFAP** | Rabbit | Astrocytes (indicates lesion area) | IgG | Sigma | G9269 | 1:500 |
| **Anti- β-tubulin III** | Mouse | Neurons (indicating neuronal growth) | IgG2b | Sigma | T8660 | 1:500 |
| **Anti-GAP43** | Rabbit | Neuronal growth | IgG | Abcam | ab16053 | 1:500 |
| **Anti-5-HT** | Goat | Neuronal growth of Raphe spinal neurons | IgG | Abcam | ab66047 | 1:500 |
| **Anti-IbA1** | Goat | Microglia (indicating neuroinflammation) | IgG | Abcam | ab5076 | 1:500 |

**D**

| Antibody | Host | Target | Supplier | Cat # | Dilution | Excitation/ Emission |
|---|---|---|---|---|---|---|
| **Alexa Fluor™ 647** | Donkey | Anti-Mouse IgG | Thermo Fisher | A-31571 | 1:500 | 650/671 nm |
| **Alexa Fluor™ 594** | Donkey | Anti-Rabbit IgG | Thermo Fisher | A-21207 | 1:500 | 590/618 nm |
| **Alexa Fluor™ 488** | Donkey | Anti-Goat | Thermo Fisher | A-11055 | 1:500 | 499/520 nm |

**Supplementary Figure 8:** Spinal cords from each group were sectioned and stained with antibodies to measure the degree of neuroinflammation and regeneration. (**A**) 20 μm thick sagittal sections were cut from 6 regions parallel to the midline of the 1 cm blocks of tissue and distributed onto slides. (**B**) Binary image of one channel/marker is shown in B1, background subtraction was performed in B2, threshold was applied in B3 to create a mask of fluorescence shown in B4. The area of fluorescently labeled tissue was then normalized against Hoechst 33342-positive cells to account for variations in tissue size. (**C**) The specific primary antibodies, their targets and dilutions are listed. (**D**) The three specific secondary antibodies that were used are listed, as well as their dilutions and excitation / emission rates.

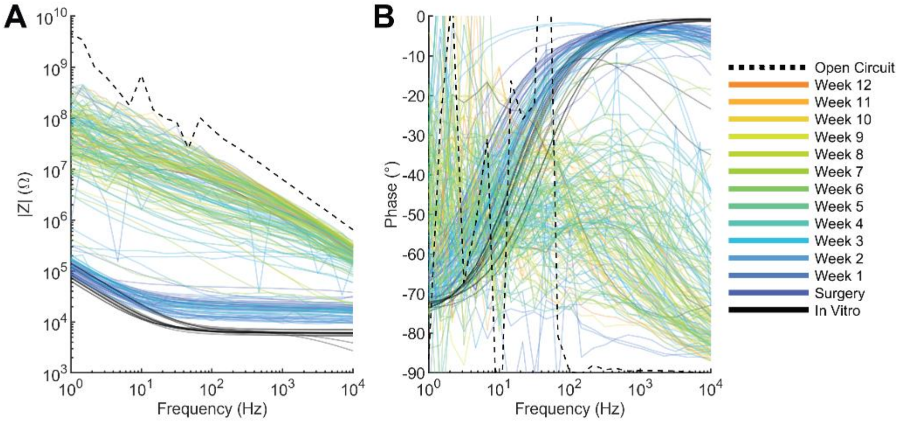


**Supplementary Figure 9:** (**A**) Impedance magnitude and (**B**) phase for reserve electrodes used for stimulation are shown over 12 weeks, indicated by the color scale, with the dashed red line representing the open circuit level.

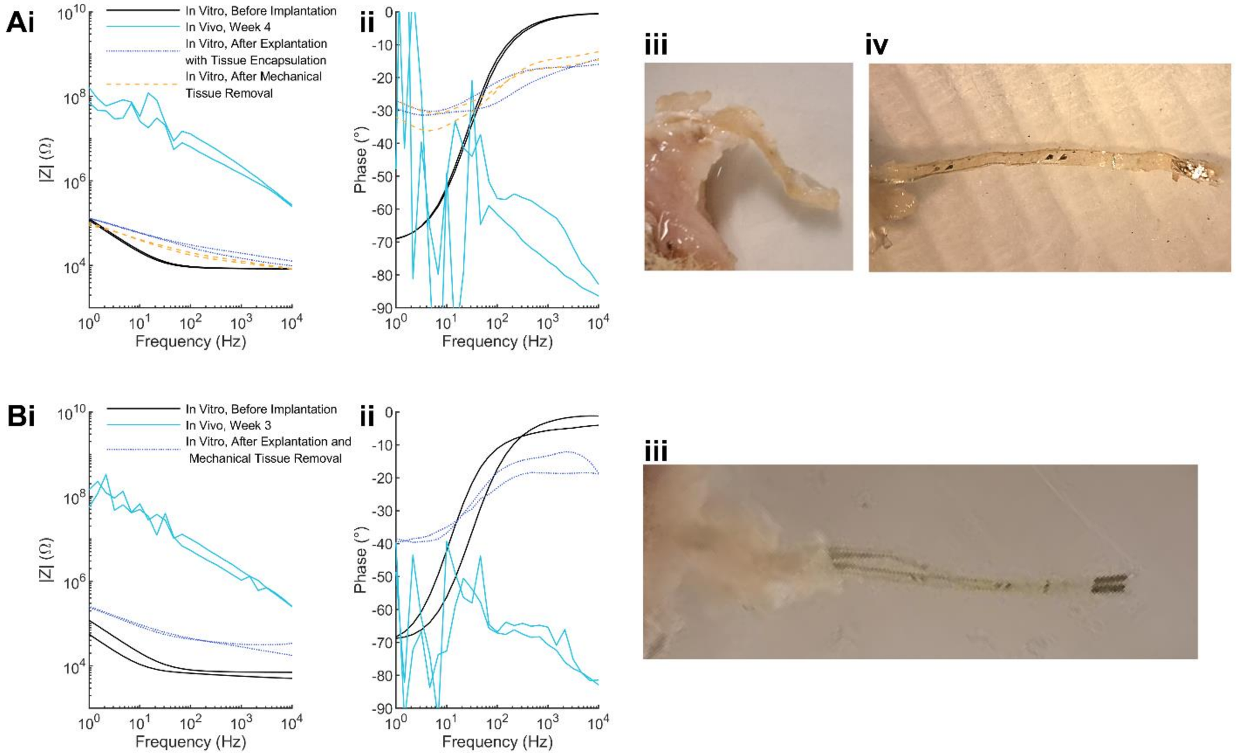


**Supplementary Figure 10:** Explants that were successfully recovered at **(A)** Week 4, and **(B)** Week 3. **(i-ii)** Impedance at various stages of examination. **(iii-iv)** Implants before and after removal of encapsulating tissue.
